# Ecotin protects *Salmonella* Typhimurium against the microbicidal activity of host proteases

**DOI:** 10.1101/2024.05.15.594389

**Authors:** L. Saposnik, L.M. Coria, L. Bruno, F.F. Guaimas, J. Pandolfi, M. Pol, M.E. Urga, F. Sabbione, M. McClelland, A. Trevani, K.A. Pasquevich, J. Cassataro

## Abstract

*Salmonella enterica* serovar Typhimurium causes acute diarrhea upon oral infection in humans. The harsh and proteolytic environment found in the gastrointestinal tract is the first obstacle that these bacteria face after infection. However, the mechanisms that allow *Salmonella* to survive the hostile conditions of the gut are poorly understood. The *ecotin* gene is found in an extensive range of known phyla of bacteria and it encodes a protein that has been shown to inhibit serine proteases. Thus, in the present work we studied the role of *ecotin* of *Salmonella* Typhimurium in host-pathogen interactions. We found that *Salmonella* Typhimurium Δ*ecotin* strain exhibited lower inflammation in a murine model of *Salmonella* induced colitis. The Δ*ecotin* mutant was more susceptible to the action of pancreatin and purified pancreatic elastase. In addition, the lack of *ecotin* led to impaired adhesion to Caco-2 and HT-29 cell lines, related to the proteolytic activity of brush border enzymes. Besides, Δ*ecotin* showed higher susceptibility to lysosomal proteolytic content and intracellular replication defects in macrophages. In addition, we found Ecotin to have a crucial role in *Salmonella* against the microbicide action of granules released and neutrophil extracellular traps from human polymorphonuclear leukocytes. Thus, the work presented here highlights the importance of *ecotin* in *Salmonella* as countermeasures against the host proteolytic defense system.

**IMPORTANCE:** The gastrointestinal tract is a very complex and harsh environment. *Salmonella* is a successful food borne pathogen, but little is known about its capacity to survive against the proteolysis of the gut lumen and intracellular proteases. Here, we show that Ecotin, a serine protease inhibitor, plays an important role in protecting *Salmonella* against proteases present at different sites encountered during oral infection. Our results indicate that Ecotin is an important virulence factor in *Salmonella*, adding another tool to the wide range of features this pathogen uses during oral infection.

## INTRODUCTION

*Salmonella* pathovars infect a wide variety of hosts and are associated to gastrointestinal or systemic diseases in humans [1, 2]. Their success relies in their ability to fight the immune system and highjack the infected cells to persist and spread to other individuals [3–5]. According to the World Health Organization, non- typhoidal *Salmonella* serovars are among the leading causes of gastrointestinal diseases around the world [6, 7] with *Salmonella enterica* serovar Typhimurium (STm) as a prevalent representative of this group.

As a food-borne pathogen, the main route of entrance of *Salmonella* is through the oral cavity, where it faces a harsh environment as it travels down the gastrointestinal tract. The changes in pH and the activity of gut proteases are not only directed to digest ingested food but also to degrade invading microorganisms in a non-specific fashion [8, 9]. It has been shown that *Salmonella* has several mechanisms to survive under low pH conditions in the stomach [10] and in the *Salmonella* containing vacuole (SCV) of infected macrophages [11]. However, little is known about how it survives against the action of gastrointestinal and intracellular proteases. *Salmonella* may synthesize protease inhibitors to hinder the proteolytic defense systems of the host and evade its responses against infections.

Ecotin is a protease inhibitor present in the genome of many gram negative bacteria [12]. Its ability to inhibit a wide range of serine proteases was proven in *Escherichia coli* [13]. It can also protect *Escherichia coli* and *Pseudomonas aeruginosa* from neutrophil elastase [14, 15]. In STm the *ecotin* gene is upregulated under stress conditions [16] and helps *Salmonella* Enteritidis to survive egg white microbicide activity [17]. Despite all these findings, the possible relevance of Ecotin in the different environments that *Salmonella* confronts during infection remains unexplored. Given that proteases are found not only in the lumen of the gastrointestinal tract but also as part of the microbicide mechanisms used by the immune system to control pathogens [18–20], we speculated that Ecotin could protect bacteria in one or both situations. Thus, we studied if Ecotin allows STm to better survive through the hostile environment of the gastrointestinal tract and against macrophages and polymorphonuclear leukocytes, which are prepared to clear a potential infection.

## RESULTS

### Ecotin has a role in STm induced colitis

To investigate the role of Ecotin in STm infection, the knock-out strain, Δ*ecotin* (Δ*eco*) and the complemented strain (Δ*eco*+*eco*) were constructed in the STm 14028s wild type (WT) background. The identity of these strains was confirmed by PCR and western blot (Fig. S1A-B). The *ecotin* deletion did not affect membrane permeability and bacterial growth since no differences were observed in N-phenyl- 1-napthylamine (NPN) uptake assays or bacterial growth curves between Δ*eco* and WT strains (Fig. S1C-D). To elucidate the role of *ecotin* during the gastrointestinal colonization, a mouse model of *Salmonella* induced colitis was used [21]. In this model, the STm infection leads to the reduction of the cecum weight, the recruitment of polymorphonuclear leukocytes (PMNs) that transmigrate from the lamina propria to the gut lumen, the reduction in the number of Goblet cells with secretory vesicles and the loss of epithelial integrity. Mice infected with WT strain had a significant reduction of cecum weight at 24h post infection (p.i.) in comparison to mock-infected mice. However, this reduction was not evidenced upon infection with Δ*eco* strain while the Δ*eco+eco* strain rescued the WT phenotype (Fig. 1A). Histopathological analysis was performed blinded (Fig. 1B) and revealed that while the WT and Δ*eco+eco* strains led to an increase in submucosa width (pronounced edema), mice infected with Δeco showed no differences compared to mock infected mice (Fig. 1C). The three STm strains induced PMN infiltration (Fig. 1D). A significant reduction of Goblet cells was exerted by WT and Δ*eco+eco* but Δ*eco* mutant failed to induce this reduction compared to mock infected mice (Fig. 1E). The loss of epithelial integrity was analyzed, with a higher score indicating more damage suffered by the epithelium. Mice infected with the Δ*eco* strain showed no differences in the score compared to the mock-infected animals, while the WT infected animals showed a higher score than mock-infected animals (Fig. 1F). The Δ*eco*+*eco* infected mice did not revert this effect. Altogether, these features revealed that Δ*eco* infection elicited a lower inflammation than the WT strain and Δ*eco*+*eco* rescued this attenuated phenotype (Fig. 1G). This lower inflammation could be due to a reduced number of Δ*eco* cells arriving at the gut epithelium. To test this hypothesis, mice were infected with WT or Δ*eco* strains and 24h p.i. the bacterial counts were determined in both the cecal contents and the cecal epithelium. Total bacterial counts in the cecal content of Δ*eco* strain were ∼10-fold lower than those of the WT strain (Fig. 1H), whereas bacteria inside host cells, which consist mainly of PMNs [22] were similar in Δ*eco* or WT infected mice. In addition, ∼10-fold lower bacterial counts were found inside the cecal tissues of Δ*eco* infected mice than in those of WT infected mice (Fig. 1I). Together, these data indicate that Ecotin in STm is important to induce colitis. Lack of *ecotin* resulted in a reduction of pathogenicity that may be related to diminished bacterial counts in cecal content and decreased invasion of the gut epithelium.

**Figure 1.**
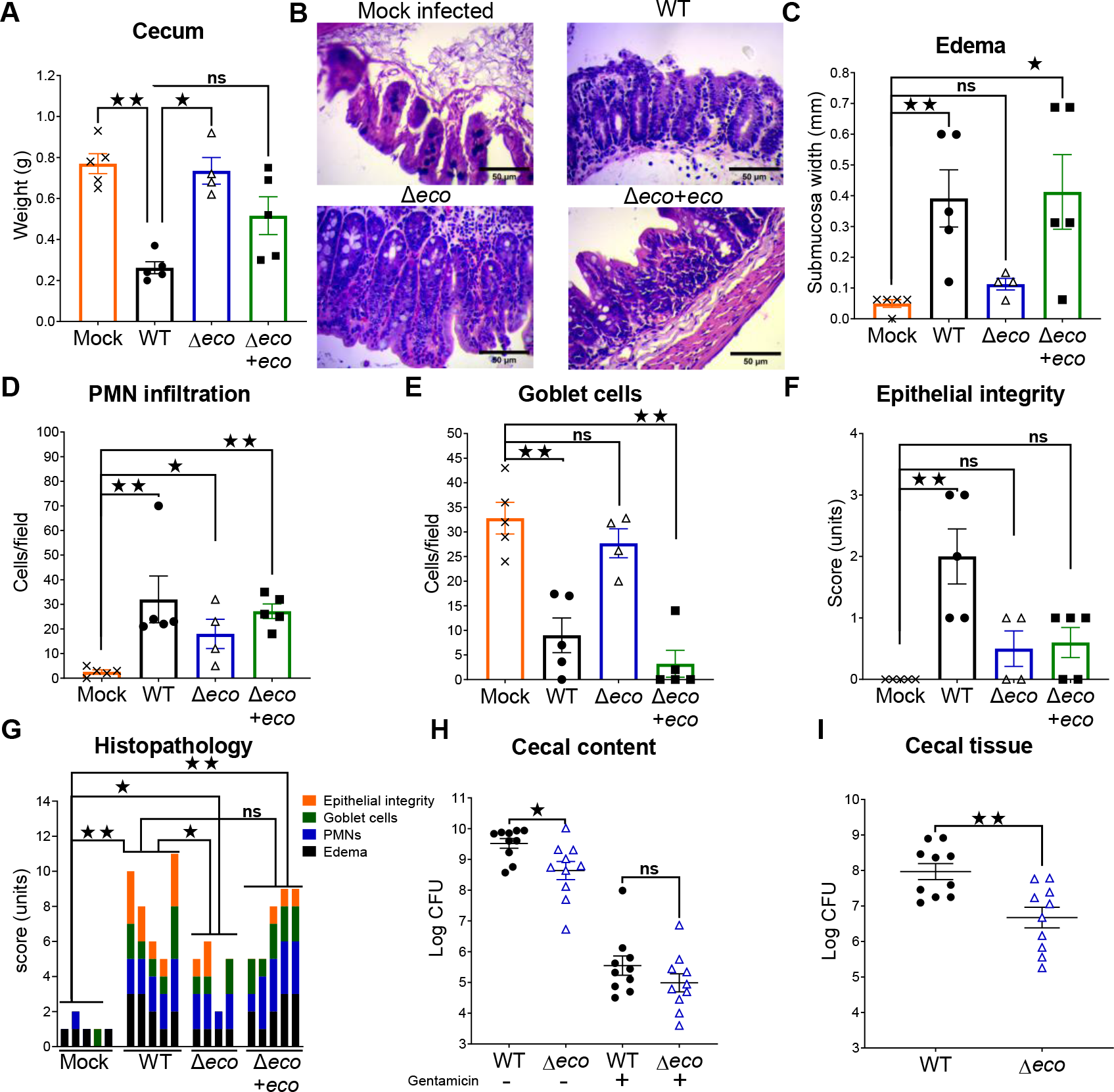
Reduced inflammation in the streptomycin pre-treatment murine infection model induced by Δ*ecotin* is associated with lower bacterial counts in the cecal epithelium. Streptomycin pre-treated mice were orally infected with 10^7^ CFU of WT (n=5), Δ*eco* (n=4), Δ*eco+eco* (n=5) strains, or mock infected with phosphate buffered saline (PBS) (n=5) and 24h p.i. were sacrificed. **(A)** Cecum was weighted separately for each mouse. Bars represent the mean±SEM. Each dot represents one mouse. Kruskal-Wallis test. **(B)** Histopathological sections of hematoxylin and eosin stained cecum tissue. Representative images of each group are shown. Bar scale 50 µm. Histopathological parameters were scored: **(C)** edema of the submucosa, **(D)** infiltration of PMNs, **(E)** amount of Goblet cells with secretory vesicles, and **(F)** epithelial integrity. Bars represent mean±SEM. Each dot represents one mouse. **(G)** Combined score. Each bar represents an individual mouse. Mann- Whitney test. Data of one representative experiment from two independent experiments. **(H-I)** Streptomycin pre-treated mice were infected with 10^7^ CFU of WT (n=10) or Δ*eco* (n=10) and 24h p.i. mice were sacrificed. Cecal contents and cecal tissue were obtained. **(H)** A fraction of the cecal content was plated, and the other half incubated with gentamicin to kill extracellular bacteria, then bacterial counts were determined. The mean±SEM is shown. Mann-Whitney test. **(I)** Bacterial counts were determined in cecal tissue. The mean±SEM is shown. Pooled data from two independent experiments. Each dot represents one mouse. Mann-Whitney test. *^ns^p>0.05*, *\*p<0.05*, *\*\*p< 0.01*.

### Ecotin protects STm against the action of pancreatic proteases

To investigate whether Ecotin is relevant for STm survival and replication in the presence of stomach proteases, WT and Δ*eco* strains were incubated *in vitro* with purified pepsin from porcine gastric mucosa or buffer (pH=3). No differences in the relative bacterial count were found between the WT and Δ*eco* strains either in buffer or with the addition of pepsin (Fig. 2A). This result indicates that Ecotin is dispensable for STm survival during pepsin exposure while travelling through the stomach. After leaving the stomach, enteric pathogens reach the intestine where they encounter pancreatic proteases. The indicated strains were incubated with buffer (pH=7) or pancreatin. Incubation with buffer alone showed no differences between WT, Δ*eco* and Δ*eco*+*eco* strains, while the addition of pancreatin resulted in a significant ∼50% (2h) and ∼75% (4h) reduction of the relative bacterial count of Δ*eco* compared to WT strain (Fig. 2B). This increased susceptibility of Δ*eco* reverted by Δ*eco*+*eco*. The susceptibility of Δ*eco* to pancreatic proteases was further studied using purified porcine pancreatic elastase, which is present in pancreatin. Incubation with buffer (pH=8.8) showed no differences between the strains, while the addition of pancreatic elastase led to significant ∼35% (2h) and ∼40% (4h) reductions of the relative bacterial count for Δ*eco* compared to the WT strain (Fig. 2C). Importantly, the Δ*eco*+*eco* rescued the observed phenotype. In addition, bacterial growth was monitored during buffer or pancreatic elastase incubation using a microplate reader. The addition of pancreatic elastase did not affect the WT strain growth curves, while the Δ*eco* strain showed a lower *plateau* in the stationary phase after pancreatic elastase incubation in comparison to the buffer condition (Fig. 2D). Evaluation of the relative carrying capacity, a parameter related to the bacterial counts reached in stationary phase, revealed that addition of pancreatic elastase led to a significant ∼15% reduction in the carrying capacity in Δ*eco* compared to the WT strain (Fig. 2D). As the indicated strains growth in LB and LB-NaCl 0.3M was similar (Figures. S1D and S2D), the lower OD620nm reached by Δ*eco* in the curves with pancreatic elastase indicated an antimicrobial effect of this protease. Taken altogether, these results indicate that expression of Ecotin in STm plays a protective role against proteases of the gut lumen.

**Figure 2.**
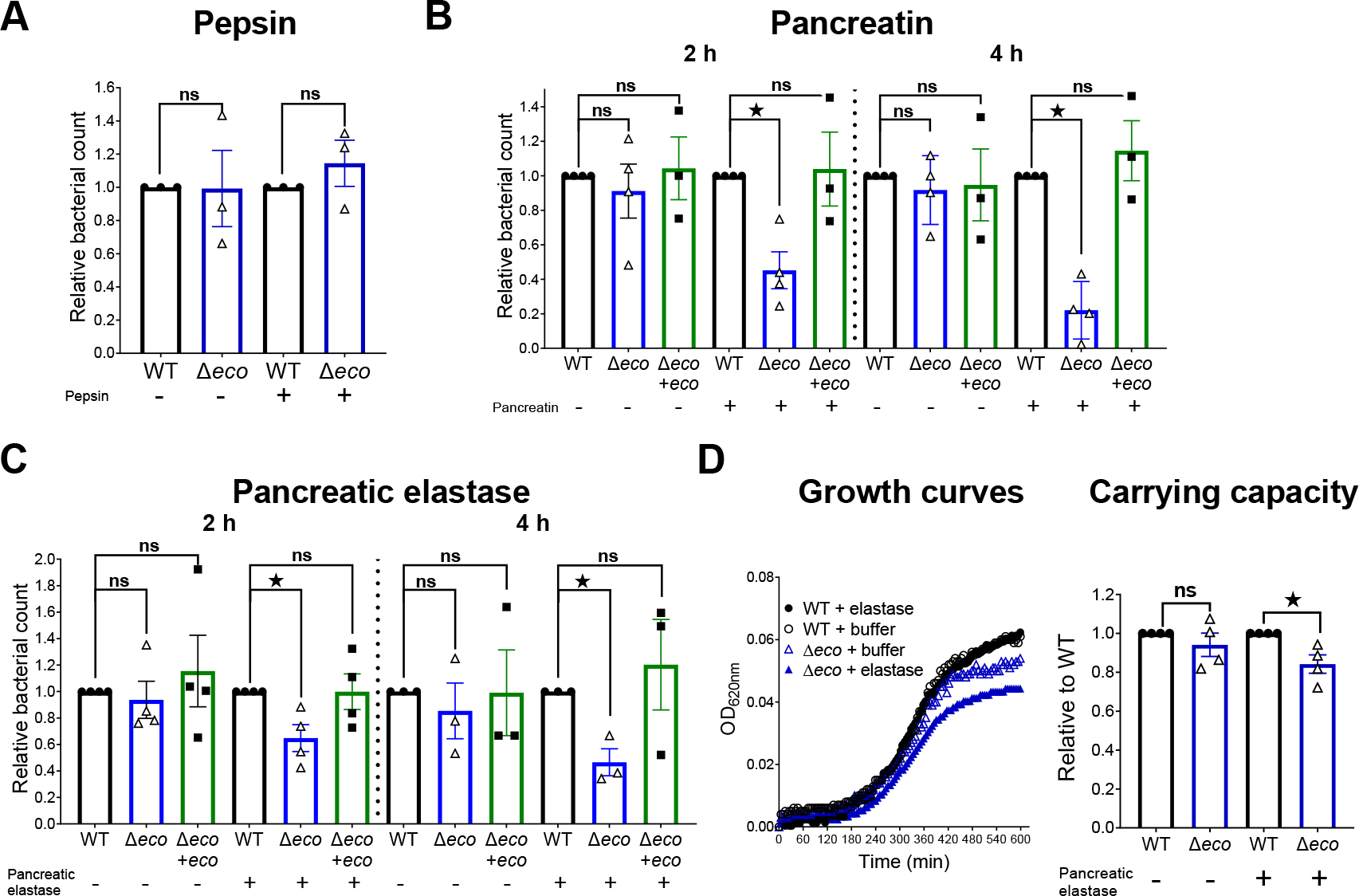
*Ecotin* allows STm to survive against pancreatic proteases. WT, Δ*eco* and Δ*eco*+*eco* were incubated with **(A)** pepsin or buffer (pH=3), **(B)** pancreatin (a complex mixture of enzymes from porcine pancreas) or buffer (pH=7) and **(C)** pancreatic elastase or buffer (pH=8.8). The bacterial counts at each time point were normalized to the WT strain (relative bacterial count). Bars represent the mean±SEM. Dots represent independent experiments. Mann-Whitney test for (A) or Kruskal-Wallis test (B-C). *^ns^p>0.05*, *\*p<0.05*. **(D)** WT and Δ*eco* strains were incubated with pancreatic elastase or with buffer alone, both conditions supplemented with LB and growth was monitored using a microplate reader. The growth curves were plotted and the carrying capacity was calculated and normalized by the WT strain. Bars represent mean±SEM. Dots represent independent experiments. Mann-Whitney test. *^ns^p>0.05*, *\*p<0.05*.

### STm Δ*ecotin* is more susceptible to the proteolytic activity of brush border membranes and has impaired adhesion to human intestinal cell lines

Upon arriving to the gut, *Salmonella* must overcome the intestinal epithelial barrier where it may confront secreted brush border proteases. Thus, Ecotin capacity to inhibit proteolytic activity of brush border membranes (BBM) from mouse intestines was tested using casein-BODIPY, a substrate that yields a fluorescent product upon digestion. BBM caused fluorescence to rapidly increase due to casein-BODIPY digestion. In contrast, BBM in the presence of recombinant purified Ecotin (rEcotin) from STm or a commercial protease inhibitor cocktail (PIC) reduced the rate of fluorescence accumulation in a dose dependent manner (Fig. 3A). Adhesion assays using HT-29 or Caco-2 human intestinal cell lines were performed. These cell lines resemble some aspects of the small intestine, the first with characteristics of a mucus secreting cell and the second being a more absorptive type of cell [23]. The adhesion of Δ*eco* strain to HT-29 or Caco-2 monolayers were ∼60% and ∼75% lower than the WT strain respectively, while Δ*eco*+*eco* rescued this phenotype in both cases (Fig. 3B). The adhesion defect of Δ*eco* was confirmed by confocal microscopy (Fig. 3C) as there were significantly fewer Δ*eco* bacteria adhered to Caco-2 monolayers than the WT strain (Fig. 3D). Moreover, in competitive experiments, bacterial adhesion was lower for Δ*eco* in both cell lines compared to WT strain (Fig. 3E), whereas the Δ*eco*+*eco* strain reverted the phenotype. Together, these data suggest that the absence of Ecotin in STm results in a loss of fitness for adhesion to the intestinal epithelium. This impaired adhesion was not due to differences in bacterial swimming and arrival to the monolayers as these strains had no differences in motility (Fig. S1E). In addition, a centrifugation step to force the bacteria onto the monolayers led to an increase in total adhesion but maintained the attenuated phenotype for Δ*eco* (Fig. S2A-B). Also, Triton X-100 used to lyse the monolayers before dilution and plating did not affect bacterial counts (Fig. S2E). These results indicated that the attenuated phenotype observed for the Δ*eco* strain is at least partially attributable to a deficient interaction of bacteria with epithelial cells because of the protease inhibitor activity on BBM exerted by Ecotin. To test this hypothesis, adhesion assays were performed pre-incubating Caco-2 monolayers with rEcotin or PIC. Incubation of monolayers with rEcotin did not affect the WT strain adhesion index, while pre-incubation of cells with rEcotin or with PIC restored the adhesion index of Δ*eco* mutant to the WT levels (Fig. 3F). These data suggest that Ecotin is important for *Salmonella* in the process of adhesion to intestinal epithelial cells.

**Figure 3.**
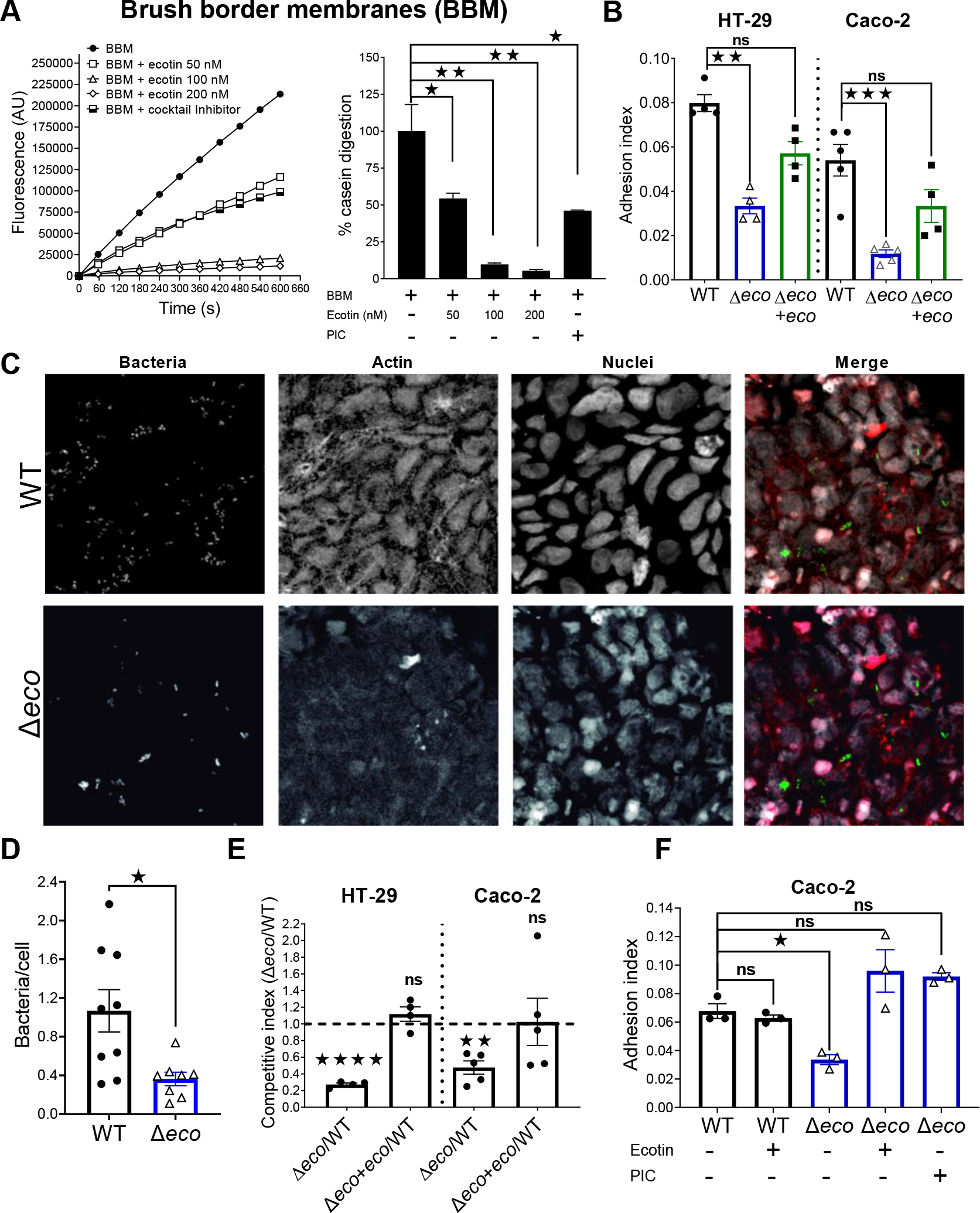
Impaired adhesion of the Δ*ecotin* strain to Caco-2 and HT-29 to epithelial cell monolayers. (A) Purified brush border membranes were co-incubated with casein-BODIPY (fluorogenic substrate) and buffer, different concentrations of recombinant Ecotin or PIC (positive control). Casein digestion was followed by fluorescence determination. The % casein digestion was calculated as percentage of the slopes of Fluorescence over time curves for each treatment compared with the buffer alone condition (100% casein digestion). Bars represent the mean±SEM. Student’s t-test. *\*p<0.05*, *\*\*p<0.01*. **(B)** Adhesion assay in HT-29 or Caco-2 cell line monolayers. Cells were incubated with WT, Δ*eco* and Δ*eco+eco* strains for 30min, washed, bacterial count determined and normalized to inoculum (adhesion index). Bars represent the mean±SEM. Dots are technical replicates. Representative of three independent experiments. Kruskal-Wallis test. *^ns^p>0.05, **p< 0.01, ***p<0.001*. **(C-D)** Adhesion assay: Caco-2 cell line monolayers were incubated for 30min with WT or Δ*eco* strains carrying GFP expressing plasmid (green). Actin filaments were stained with phalloidin (red) and nucleus were stained with Topro-3 (white). Bacteria in random fields were counted and normalized by the number of nuclei per field. Bars represent the mean±SEM. Dots are random fields. Pooled data from two independent experiments. Mann-Whitney test. *\*p<0.05*. **(E)** Competitive adhesion assay performed in HT-29 or Caco-2 cell line monolayers. The competitive index was calculated as the ratio in bacterial count after 30min of infection corrected by the ratio in the inoculum. Bars represent mean±SEM. Dots are technical replicates. Representative of at least two independent experiments. One Sample t-test vs 1. *^ns^p>0.05*, *\*\*p< 0.01*, *\*\*\*\*p<0.0001*. **(F)** Adhesion assays in Caco- 2 were repeated with the pre-treatment of the indicated wells with recombinant Ecotin 1 µM or PIC for 5min before the addition of bacteria. Bars represent the mean±SEM. Dots are technical replicates. Representative of two independent experiments. One-Way ANOVA. *^ns^p>0.05*, *\*p<0.05*.

### STm Δ*ecotin* is more susceptible to lysosomal proteolytic content and has impaired intracellular replication in murine macrophages

STm infects macrophages, which allow bacteria to disseminate to other tissues. To study the possible contribution of Ecotin to bacterial intracellular survival and replication in macrophages, the murine macrophage J774 cell line was infected with the different STm strains. There were no differences in invasion capacity between the strains (Fig. 4A). However, the Δ*eco* mutant had a significantly ∼50% reduction in intracellular replication after 4h p.i. compared to the WT strain and this phenotype was rescued by Δ*eco*+*eco* complementation (Fig. 4B). The contribution of Ecotin to intracellular replication was also demonstrated in competitive assays, no differences in invasion between Δ*eco* and WT strain were observed, while a significant fitness advantage for WT strain over the Δ*eco* mutant was observed at 4h p.i. (Fig. 4C). To evaluate if the observed results were linked to differences in growth rates, the TIMER^bac^ plasmid was used as a growth rate reporter of *Salmonella* [24]. Thus, J774 macrophages were infected with WT or Δ*eco* strains carrying TIMER^bac^ and confocal microscopy analysis was performed. There were no differences in green/orange fluorescence ratio between Δ*eco* and WT strains at invasion but the Δ*eco* mutant showed a significantly lower ratio at 4h p.i. than the WT strain (Fig. 4D). These results suggest that Δ*eco* intracellular growth is limited in macrophages. To further study this phenomenon, the intracellular trafficking of Δ*eco* and WT strains was studied. Recruitment of lysosome-associated membrane protein 1 (LAMP-1), a late endosome marker, related to the maturation of the SCV [25, 26] was evaluated. The Δ*eco* mutant recruited significantly more LAMP-1 at the invasion time point than the WT strain (36%±3% vs 24%±3%), but this difference was abolished at 4h p.i., showing a distinctive intracellular traffic dynamic for Δ*eco* mutant (Fig. 4E-F). To assess the role of Ecotin against the lysosomal proteolytic content, J774 derived microsomes with demonstrated proteolytic activity (Fig. S3A) were incubated with the Δ*eco* mutant or the WT strain. There were no differences between Δ*eco* and WT strains when incubated with buffer, but notably a ∼40% reduction of relative bacterial counts for Δ*eco* mutant was observed compared to the WT strain after incubation with microsomes (Fig. 4G). In this context, the lower intracellular replication found at 4h p.i. could be due to an impaired ability of the Δ*eco* mutant to survive against the action of intracellular proteases. These results show that Ecotin is an important virulence factor of STm for establishing a replicative niche in macrophages.

**Figure 4.**
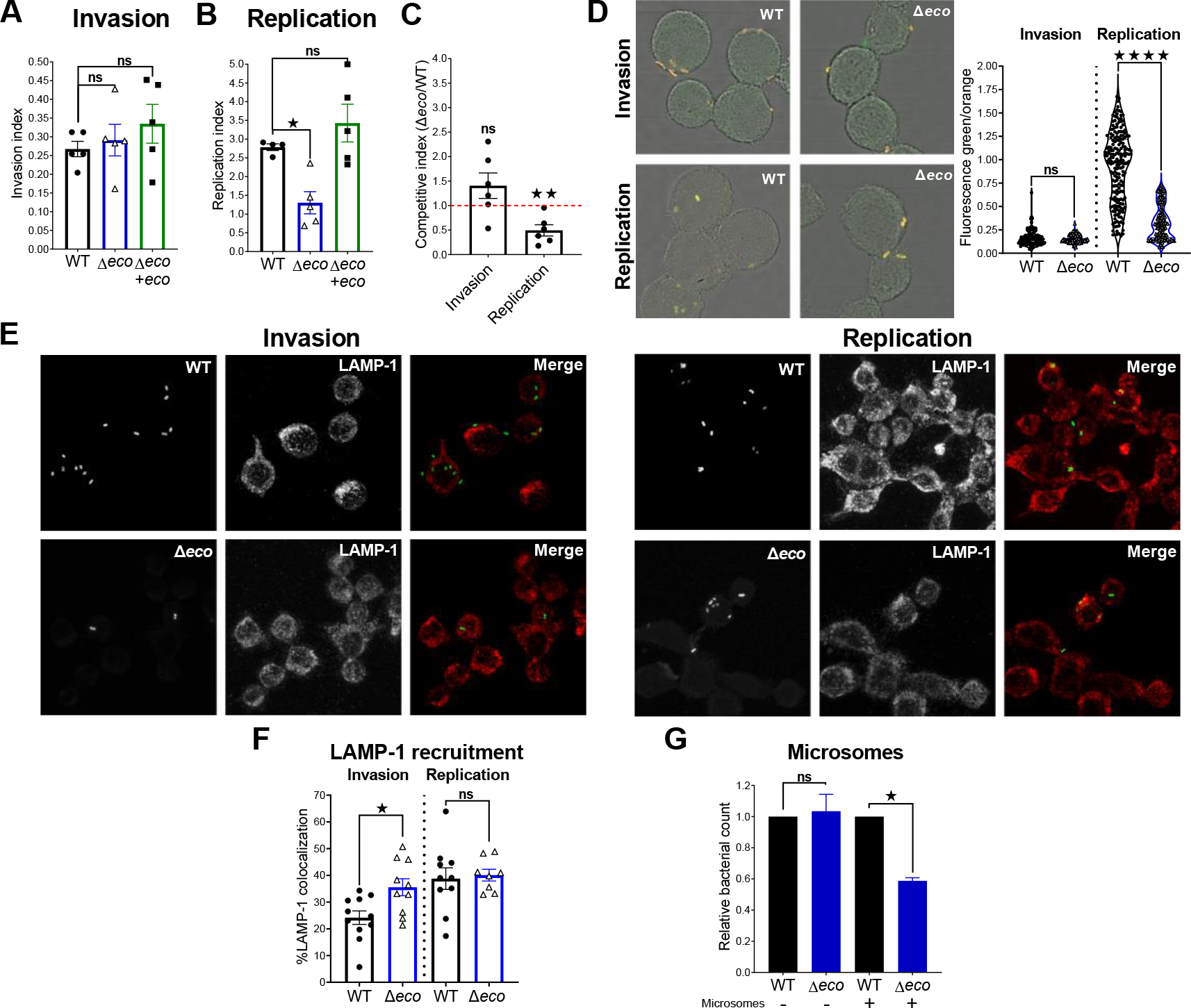
Δ*ecotin* has an attenuated intracellular replication in murine macrophages and is more susceptible to the action of intracellular proteases. **(A)** J774 macrophages were incubated with WT, Δ*eco* or Δ*eco*+*eco* strains for 30min. Then, cells were treated with gentamicin for 1h, washed, lysed and bacterial counts determined. Bacterial counts were normalized by the inoculum (invasion index). Bars represent the mean±SEM. Dots are technical replicates. Representative of three independent experiments. One-Way ANOVA with Bonferroni post-test. *^ns^p>0.05*. **(B)** Cells were infected as in A, but incubated for a further 3h before washing and bacterial counts determination. The bacterial counts at this time point were then normalized by the bacterial counts in the invasion time point (replication index). Bars represent the mean±SEM. Dots are technical replicates. Representative of three independent experiments. One-Way ANOVA with Bonferroni post-test. *^ns^p>0.05*, *\*\*p< 0.01*. **(C)** Competitive assay performed in J774 macrophages. The competitive index was calculated as the ratio between Δ*eco*/WT recovered at the invasion time point or 4h time point corrected by inoculum or invasion ratio respectively. Bars represent the mean±SEM, Dots are technical replicates. Representative of three independent experiments. One-Sample t-test vs 1. *^ns^p>0.05*, *\*\*p< 0.01*. **(D)** J774 macrophages were infected with WT or Δ*eco* strains carrying a TIMER^bac^ plasmid and fluorescence intensity analyzed by confocal microscopy. The ratio of green/orange fluorescence of TIMER^bac^ indicates the growth rate of *Salmonella*, slow-growing cells appear orange/red and faster growing cells are greener. Bars represent the mean±SEM. Dots represent individual bacteria from at least 10 random fields. Pooled data from two experiments. Mann-Whitney test. *^ns^p>0.05*, *\*\*\*\*p< 0.0001*. **(E-F)** J774 macrophages were infected with WT or Δ*eco* strains expressing GFP, then cells were fixed and labeled with a primary anti LAMP- 1 antibody and a secondary antibody coupled with AlexaFluor647. Colocalization was analyzed in random fields using Manders approach to measure LAMP-1 recruitment. Bars represent the mean±SEM. Dots are random fields pooled from two experiments. Student’s t-test. *^ns^p>0.05*, *\*p< 0.05*. **(G)** Microsomes from J774 macrophages or buffer were co-incubated for 2h with WT or Δ*eco* strains. The bacterial count was normalized by WT strain. Bars represent the mean±SEM. Pooled data from two experiments. Student’s t-test. *^ns^p>0.05*, *\*p< 0.05*.

### Ecotin protects STm from the proteolytic microbicide activity of human PMNs

PMNs recruitment is a hallmark of STm induced colitis and these cells are the main immune system cells that fights and clears *Salmonella* infection [27]. As it was shown that Ecotin from *Escherichia coli* and *Pseudomonas aeruginosa* was able to inhibit neutrophil elastase [14, 15] we wanted to corroborate the inhibitory capacity of STm Ecotin against it. Using recombinant Ecotin, we confirmed that it could inhibit neutrophil elastase activity in a dose dependent manner (Fig. 5A). To test if Ecotin could be involved in bacterial survival against PMN proteolytic microbicide activity, purified human PMNs were co-incubated with the different *Salmonella* strains. Human PMNs were able to similarly kill WT, Δ*eco* and Δ*eco*+eco strains resulting in a ∼50% survival. However, when the oxidative dependent microbicide activity was inhibited by adding DPI to PMNs prior to infection, this treatment revealed that the Δ*eco* strain was more susceptible than the WT strain to the non-oxidative killing mechanisms of PMNs (Fig. 5B). This effect was reverted by Δ*eco*+*eco.* In addition, the Δ*eco* strain was more susceptible than the WT strain to the killing activity of supernatants of degranulated PMNs, with a ∼40% reduction in survival (Fig. 5C), or to the killing by purified neutrophil extracellular traps (NETs), with roughly ∼50% reduction in survival compared to WT strain (Fig. 5D), both with confirmed neutrophil elastase activity (Fig. S4A-B) and DNA content in the case of NETs (Fig. S4C). The Δ*eco*+*eco* strain reverted the effect of NETs to WT levels and remarkably, the addition of rEcotin to the NETs restored the survival of Δ*eco* strain to the WT levels (Fig. 5D). These results highlight the importance of Ecotin for STm survival to the PMNs proteolytic microbicide activity.

**Figure 5.**
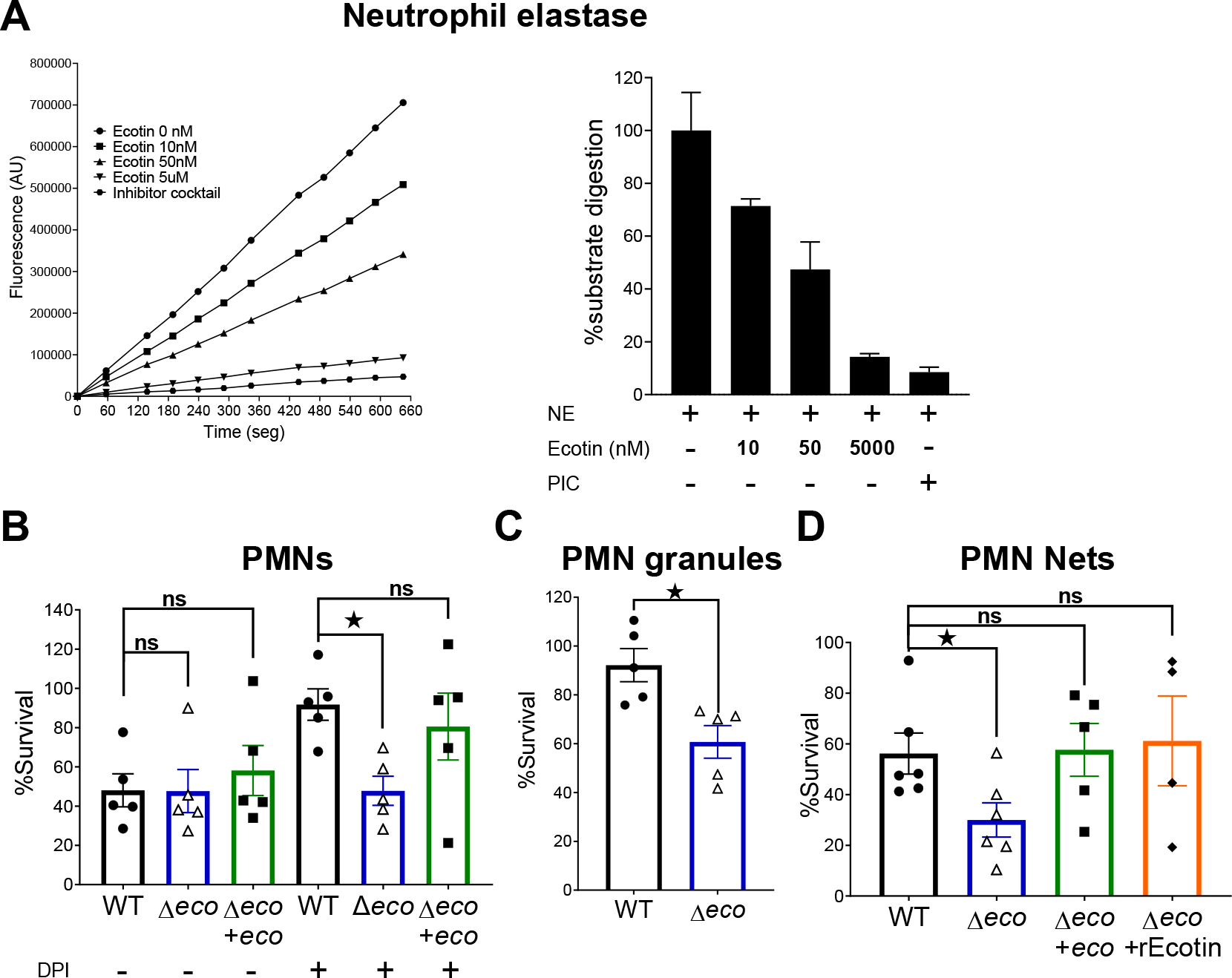
Ecotin protects STm from the proteolytic microbicide activity of human PMNs. (A) Neutrophil elastase (NE) was incubated with a fluorogenic substrate and different concentrations of recombinant Ecotin or a commercial protease inhibitor cocktail (PIC). Protease activity was measured over time using a fluorescence microplate reader. The percentage of substrate digestion was calculated as the percentage of fluorescence vs. time slope regarding the condition without inhibitor (0nM) (100% substrate digestion). **(B)** Purified human PMNs were incubated with WT, Δ*eco* or Δ*eco*+*eco* strains in the presence of absence of DPI and after 90min bacterial counts were determined and normalized to bacteria incubated without neutrophils (100% Survival). Bars represent the mean±SEM. Dots are individual donors. One-Way ANOVA with Bonferroni post-test. *^ns^p>0.05*, *\*p<0.05*. **(C)** Degranulation was induced in purified human PMNs and the supernatant containing the granules was incubated with WT or Δ*eco* strains. After 90min of incubation the bacterial counts were determined and normalized to bacteria incubated without granules (100% Survival). Bars represent the mean±SEM. Dots are individual donors. Student’s t-test. *\*p<0.05*. **(D)** Neutrophil extracellular traps (NETs) were induced in purified human PMNs and the supernatant containing the NETs was incubated with WT, Δ*eco,* Δ*eco*+*eco* strains or Δ*eco* supplemented with recombinant Ecotin (Δ*eco*+rEcotin). Then, 90min after incubation, the bacterial counts were determined and normalized to bacteria incubated without NETs (100% Survival). Bars represent the mean±SEM. Dots are individual donors. One-Way ANOVA with Bonferroni post-test. *^ns^p>0.05*, *\*p<0.05*.

## DISCUSSION

STm infection in humans causes a diarrheal illness characterized by acute intestinal inflammation and the presence of neutrophil infiltration [28]. In mice, the streptomycin pre-treated mouse model resembles many features of human gastroenteritis [21]. The inflammation induced by STm, far from killing the pathogen, creates a suitable niche for bacterial replication [29–31]. Our findings indicate that *ecotin* plays a role in *Salmonella* induced gastrointestinal inflammation, since STm lacking the *ecotin* gene induced an attenuated inflammation in infected mice.

Recruitment of PMNs into and across the gut mucosa is a cardinal feature of intestinal inflammation in response to STm infection and it has been shown that intraluminal PMNs protect the cecal epithelium from STm induced loss of integrity [32]. Here we found that Ecotin was dispensable for PMN recruitment and bacteria amount inside the luminal PMNs were similar upon WT or Δ*eco* STm infection. The reduced inflammation observed could be related to fewer bacterial counts of Δ*eco* in either the cecum lumen or invading the epithelium.

In previous studies, it was shown that bacterial protease inhibitors were important for *in vitro* survival as well as to establish infection *in vivo* [33]. The STm Ecotin protein is exported to the periplasm via the signal peptide in its sequence (Fig. S5A). Given its periplasmatic location, Ecotin may be helping STm to defend against proteases present in the gastrointestinal tract.

We demonstrated that knocking out the *ecotin* gene rendered bacteria susceptible to proteolysis by pancreatin and pancreatic elastase. Similar to our findings, it was shown that Ecotin from *Campylobacter rectus* and *Campylobacter showae* rescued *Campylobacter jejuni* Δ*pglB* from proteolysis of chicken cecal contents [34]. These findings indicate that Ecotin protects *Salmonella* against the proteolytic activity of gut luminal proteases.

Another important source of proteases is the brush border of the small intestine, which degrade proteins and other nutrients in a non-specific fashion [35]. Here, we have shown the ability of Ecotin to inhibit the proteolytic activity of mice purified BBM.

After swimming and sensing the surface, *Salmonella* establishes the infection by invading epithelial cells [36, 37]. The initial adhesion is critical to allow the bacterial invasion machinery to inject effectors into the host cell cytoplasm in order to promote its own endocytosis [38]. We demonstrated that the Δ*eco* strain exhibited lower adhesion to HT-29 and Caco-2 cell lines than the WT strain. Moreover, this phenotype was rescued upon pre-incubation of cell monolayers with Ecotin or a PIC. These results suggest that Ecotin could be inhibiting at least one brush border protease whose biological activity may impair the adhesion of STm. We then propose that the intestinal brush border enzymes have a role in keeping the pathogen at bay.

Our results also demonstrated that Ecotin is required for survival and replication inside macrophages. Similar attenuated replication was found for a Δ*eco* mutant in *Burkholderia pseudomallei* in macrophages [39]. Once inside the macrophage, *Salmonella* changes its gene expression program to form the SCV and to shift to intracellular adaptation [40–42]. This shift made by *Salmonella* is within the phagolysosomal pathway, which is modulated by the bacteria to evade antimicrobial activity. The recruitment of LAMP-1 is typical of an intermediate to late stage of the SCV formation [43]. Interestingly, we found an altered dynamic of LAMP-1 recruitment, indicating a faster maturation for SCV containing the Δ*eco* mutant inside macrophages. This suggested a role for Ecotin in the SCV and phagolysosome maturation dynamics. In addition, the susceptibility of Δ*eco* to macrophage-derived microsomes could explain the hampered intracellular replication of the Δ*eco* mutant in this cell type. Macrophages try to kill and digest engulfed bacteria using a wide array of antimicrobial peptides, organic acids, proteases, low pH, and ROS [20, 44–47]. It has been demonstrated that murine macrophages impair STm intracellular replication by CRAMP, an antimicrobial peptide, in a serine-protease-dependent manner [48]. Also, Perforin-2 from murine macrophages, induce the formation of pores in the outer membrane of engulfed *Salmonella*. Thus, allowing the entrance of antimicrobial peptides and proteases to the periplasmic space of bacteria [49]. Consequently, the breach in the outer membrane takes the battlefield to the periplasm where Ecotin could protect *Salmonella*’s proteins from proteolysis of invading proteases. These results highlight the importance for *Salmonella* to have periplasmic protease inhibitors to defend against the host proteases while running against the clock to establish a replicative niche inside macrophages.

STm hides inside macrophages and epithelial cells to escape from PMNs killing activity [50]. Our experiments co-incubating STm and PMNs showed that abrogation of ROS-dependent microbicidal mechanisms by DPI practically abolished the ability of PMNs to kill the STm WT strain but not Δ*eco*. Under these circumstances, the Δ*eco* had reduced survival when co-incubated with PMNs, indicating that inhibiting proteases could be important for survival at any time when ROS does not dominate.

PMNs have different ways of killing pathogens; phagocytosis and digestion, release of granules and NETs, each of these using ROS, antimicrobial peptides or proteases as effectors of antimicrobial activity [19, 51]. NETs consist of PMN content using DNA as a trap, together with active neutrophil elastase, and myeloperoxidase [52]. Here we found that Ecotin is important for STm to survive against both PMN killing mechanisms, granules content and NETs. Similar results were found for *ecotin* from *Campylobacter rectus* and *Campylobacter showae* [34]. We also found that the addition of rEcotin to the NETs prior to the incubation with the Δ*eco* strain reverted the phenotype to WT levels. In previous work it was demonstrated that rEcotin from STm can inhibit neutrophil elastase present in purified PMNs [53]. Here we show that rEcotin from STm can inhibit purified neutrophil elastase. This enforced the idea that inhibition of proteases, presumably neutrophil elastase, in the NETs or granules augmented bacterial survival. Thus, our data indicate that Ecotin is a defense mechanism in STm against the antimicrobial activity of PMN serine proteases.

In summary, we demonstrated for the first time, the importance of a protease inhibitor in STm as a key protein for defending the bacteria against the host proteolytic defense system. Our experiments demonstrated the ability of Ecotin to inhibit proteolytic activity in different environments *Salmonella* encounters as it develops the infection.

## MATERIALS AND METHODS

### Bacterial strains and growth conditions

The STm strains, plasmids and primers used in this study are listed in Table S1. The Δ*ecotin* strain was constructed as described [54] and transduced into a clean WT background using the p22 phage (Δ*eco*). The primers EcotinFPcomple and EcotinRVcomple were used for amplification of the *ecotin* gene region for cloning in pWSK29 plasmid. Then, complementation was made by electroporation of the pWSK29 plasmid containing the *ecotin* gene (Δ*eco*+*eco*). Single colonies were grown overnight in LB (37°C, agitation). Next, diluted in LB-NaCl 0.3M and incubated further (37°C, 100rpm) to mid-exponential phase. If necessary, media or agar plates were supplemented with antibiotics (kanamycin-60µg/mL, ampicillin-100µg/mL).

### *Salmonella* colitis induced mouse model

Female BALB/c mice (6–9-week-old) were bred at IIB-UNSAM and orally infected as described previously [21]. At 24h p.i., mice were sacrificed, and ceca were aseptically collected. In some experiments, the ceca were weighted, and a section was suspended in 4% paraformaldehyde (PFA) (Sigma) for histopathological analysis by pathologists in a blinded manner according to previously described score guidelines [21]. In other experiments cecal contents were obtained, fractioned, and treated (30min, 37°C) with PBS (total bacterial counts) or gentamicin 400µg/mL (bacteria inside suspended cells). Then, fractions were centrifuged at 10000×g (10min), washed twice with PBS, and plated in agar-SS (Britannia). The cecal epithelium was opened, washed with PBS, incubated with gentamicin 100µg/mL (1h, 37°C). Then, washed twice with PBS, centrifuged 3000×g (5min), suspended in PBS, homogenized and plated in agar-SS.

### Proteases susceptibility assay

STm strains (1×10^5^ CFU) were incubated with pepsin (3mg/mL, Sigma) or buffer for 30min, pancreatin (2mg/mL, Sigma), pancreatic elastase (5µM, Sigma) or buffer (negative control) for 2h or 4h (37°C). Buffers were: NaCl 125mM, KCl 7mM, NaHCO3 45mM, pH=3 (pepsin), 0.5% NaCl (pancreatin) or Tris-HCl 10mM, pH=8.8 (pancreatic elastase). Bacterial counts were determined by serial dilutions and plating. Relative bacterial counts to the WT strain were calculated at each time point/condition. To study growth kinetics of STm strains in presence of pancreatic elastase, WT or Δ*eco* were incubated with buffer or pancreatic elastase as described above plus 9% LB in a microplate reader (Tecan) with agitation (37°C). The OD620nm was recorded at 5min intervals. Growth curves were analyzed using the GrowthCurver R package [55]. The carrying capacity, which is the maximum size a certain population can achieve in a particular environment, was obtained for each individual curve from the adjusted model used by the GrowthCurver. Then, the values were normalized by the carrying capacity of the WT strain in each experiment.

### Brush border membranes preparation and casein digestion inhibition

BBM were obtained by MgCl2 precipitation as previously described [56]. For BBM inhibition assays, BBM were incubated with buffer (PBS), Ecotin or PIC using casein- BODIPY (Life-Technologies) 1µg/mL as substrate. Substrate-cleavage led to fluorescence release that was determined (FilterMaxF5, Molecular-Devices). The slopes of linear parts of the curves were normalized by the buffer condition and the percentage of casein digestion was calculated.

### HT-29 and Caco-2 adhesion and competitive assays

Caco-2 and HT-29 were cultured in RPMI (Gibco) supplemented with 10% heat-inactivated fetal bovine serum, 1mM sodium pyruvate (sigma), 2mM L-glutamine (Sigma), 100U/mL penicillin (Sigma) and 100 µg/mL streptomycin (Sigma) as previously described [56]. Cells from passages 5-25 were used and seeded (5×10^4^ cells/well) in 24-well plates (Sarstedt) and allowed to reach polarization [57]. The last media change before experiments was done without antibiotics. Infections were performed with 1×10^7^ CFU/mL in RPMI for 30min (37°C). Then, monolayers were washed 4× using 0.9% NaCl and disrupted with 0.1% Triton X-100. Serial dilutions were performed and plated. The adhesion index was calculated as CFU/mL 30min p.i. by CFU/mL in the inoculum. In some experiments, Caco-2 monolayers were pre-incubated with rEcotin or PIC (5min) before infection. For competitive assays, Caco- 2 or HT-29 monolayers were incubated with bacteria as described above but using a 1:1 mix of WT:Δ*eco* or WT:Δ*eco*+*eco*. Bacteria were distinguished using different antibiotics. The ratios of Δ*eco*/WT and Δ*eco*+*eco*/WT after 30min of incubation were divided by the ratios in their respective inoculum to obtain the competitive index.

### Confocal microscopy of Caco-2

STm strains carrying the pNCS-mClover3 plasmid were used for adhesion assays in Caco-2 cells grown in coverslips. Monolayers were washed and fixed with 4% PFA, permeabilized and stained (PBS, 0.2% Triton X-100, 5% BSA, Alexa-Fluor488 phalloidin and Topro-3) for 30min. Then, coverslips were washed with water and mounted on glass slides using Fluorsave (Merck). Random fields were acquired using an IX-81 Olympus microscope with FV-1000 confocal module. Bacteria and Caco-2 nuclei were counted using Icy Microscopy Software, and the bacteria/nuclei ratios were calculated per field. The pNCS-mClover3 was a gift from Michael Lin (Addgene-plasmid #74236).

### J774 invasion, replication and competitive assays

J774 macrophages were cultured in RPMI as described for Caco-2. Cells (10^5^ cells/well) were allowed to attach in 24-well plates (24h) without antibiotics. Then, macrophages were washed with PBS and incubated with the different STm strains (MOI:50, 30min), washed twice and treated with gentamicin (100µg/mL, 1h). Afterwards, cells were 2x-washed and lysed with 0.1% Triton X-100 (invasion) or further incubated with gentamicin 20µg/mL for 3h, washed and lysed (4h -replication point-). Bacterial counts and inocula were determined by serial dilutions and plating. The invasion or replication index were calculated as CFU/mL at invasion time point divided by the inoculum CFU/mL or CFU/mL at 4h divided by the CFU/mL at invasion, respectively. Competitive assays were performed as described for Caco-2 but using the specific time points for J774 infections.

### Growth analysis and LAMP-1 recruitment in J774 macrophages infected with STm strains

The invasion and replication assays in macrophages were performed as described above with WT or Δ*ecotin* strains carrying the pBR322-TIMER. DS-red (dicroic longpass 640nm with filter 590/60) and green (dicroic longpass 560nm and filter 515/20) fluorescence after excitation with a 488nm laser were acquired simultaneously. The background was subtracted and the green/orange-fluorescence was plotted for each bacterium. The pBR322-TIMER was a gift from Dirk Bumann (Addgene-plasmid #103056). LAMP-1 recruitment was analyzed in infected macrophages with STm strains carrying the pNCS-mClover3. LAMP-1 was stained as described previously for LAMP-2 but using a rat-IgG anti-mouse LAMP-1 and anti-rat-IgG-AlexaFluor647 (Biolegend) [56]. Images of random fields were acquired, and the background was subtracted. Colocalization was calculated by the Manders approach for each field using the Icy Microscopy Software.

### Incubation of STm with J774 derived microsomes

The J774 microsomes were obtained as described before [58]. STm strains (1x10^4^ CFU) were incubated (2h) with or without microsomes in buffer (DTT 2mM, sodium citrate 50mM, pH=4.5) and then bacterial counts were determined. The relative bacterial counts were obtained by normalizing the CFU by WT strain in each condition.

### Neutrophil elastase inhibition

Neutrophil elastase (10nM) was incubated with buffer (Tris-HCl 10mM, pH=8.0), Ecotin or PIC (positive control) and N-Methoxysuccinyl-Ala-Ala-Pro-Val-AMC (Sigma) 20µM as substrate. Cleavage of the substrate led to fluorescence increase and was recorded (10min). The slope of the linear part of the curves was normalized by the buffer condition.

### Incubation with human PMNs

Peripheral blood was obtained from venipuncture of human healthy adult donors and then deposited in anticoagulated tubes. The PMNs were isolated by centrifugation on Ficoll-Paque, dextran sedimentation, and hypotonic lysis. Purity of PMN preparation was more than 98% and cells were used immediately after isolation. The STm strains (10^6^ CFU) were co-incubated with PMNs (10^6^ cells) in RPMI with 10% FBS (inactivated at 65°C) for 90min (37°C). DPI 10µM was added to PMNs 20min before infection. The survival was calculated as CFU/mL in presence of neutrophils over CFU/mL in medium.

### Incubation with neutrophil granules content

Supernatant of degranulated PMNs were obtained after treatment with cytochalasin-B and fMLP as previously described [59]. Then, 10^6^ CFU of the STm strains were mixed with supernatants of ∼10^6^ degranulated PMNs for 90min (37°C). The survival was calculated as CFU/mL in presence of granules over CFU/mL in medium.

### Incubation with neutrophil extracellular traps

Purified human PMNs were incubated with phorbol 12-myristate 13-acetate (PMA) (Sigma) 100nM for 4h. Then, DNaseI (Sigma) (2U/mL) was added for 10 min (37°C). Afterwards, EDTA (5mM) was used and supernatant containing NETs separated by centrifugation (3000×g). STm strains (10^6^ CFU) were mixed with the NETs of ∼10^6^ PMNs for 90 min (37°C). The survival was calculated as CFU/mL in presence of NETs over CFU/mL in culture medium. The rEcotin was added to the NETs 5min before the co-incubation with the Δ*eco* strain.

### Ethics Statement

The participation of human blood donors was reviewed and approved by Central Ethical Committee of the Buenos Aires Province (ACTA-2023-08191112-GDEBA-CECMSALGP). Written informed consent to participate in this study was provided by the participants. Protocols of this study agreed with international ethical standards for animal experimentation (Helsinki Declaration and amendments, Amsterdam Protocol of welfare and animal protection and NIH guidelines for the Care and Use of Laboratory Animals). Protocols of this study were approved by the Institutional Committee for the Care and Use of Experimentation Animals from UNSAM (CICUAE-UNSAM_N°17/2018).

### STATISTICAL ANALYSIS

Statistical analyses were performed using GraphPad Prism (GraphPad Software, Inc., San Diego, CA, USA). The normality of the data was tested to use parametric tests if possible. The statistical tests used are indicated in figure legends. A p<0.05 was considered statistically significant. ns=not statistically significant (p>0.05).

## ACKNOWLEDGMENTS

This work was supported by the National Agency of Promotion of Science and Technology (ANPCyT) from Argentina by grants PICT 2019 0625 and PICTA BCEI CAT III 2021 48 to JC. MM was supported, in part, by US National Institutes of Health grants R01 AI136520 and R03 AI139557. We thank Dr. Susana K. Checa for kindly giving us the p22 phage. We also thank Rosario Cespedes and Jose Acosta for helping in the obtention of blood samples. Finally, we thank Weiping Chu and Steffen Porwollik for technical help.

